# Interaction of replication factor Sld3 and histone acetyl transferase Esa1 alleviates gene silencing and promotes the activation of late and dormant replication origins

**DOI:** 10.1101/2020.09.21.305680

**Authors:** Seiji Tanaka

## Abstract

DNA replication in eukaryotes is a multi-step process that consists of three main reactions: helicase loading (licensing), helicase activation (firing), and nascent DNA synthesis (elongation). Although the contributions of some chromatin regulatory factors in the licensing and elongation reaction have been determined, their functions in the firing reaction remain elusive. In the budding yeast *Saccharomyces cerevisiae*, Sld3, Sld7, and Cdc45 (3-7-45) are rate-limiting in the firing reaction and simultaneous overexpression of 3-7-45 causes untimely activation of late and dormant replication origins. Here we found that 3-7-45 overexpression not only activated dormant origins in the silenced locus, *HMLα*, but also exerted an anti-silencing effect at this locus. For these, interaction between Sld3 and Esa1, a conserved histone acetyltransferase, was responsible. Moreover, the Sld3–Esa1 interaction was required for untimely activation of late origins. These results reveal the Sld3–Esa1 interaction as a novel level of regulation in the firing reaction.

## INTRODUCTION

In eukaryotes, DNA is stored in the nucleus as specific protein–nucleic acid complexes called chromosomes. Each chromosome is made up of basic units called nucleosomes, each of which contains DNA and a histone octamer. The properties of the nucleosomes are regulated by the post-translational modification of histones. Differences in these modifications generate local chromatin structure, which in turn affects transcription capacity and the efficiency of DNA repair. However, the effects of chromatin on DNA replication are not well understood.

Eukaryotic DNA replication initiates from multiple specific regions of chromosomes, called replication origins. DNA replication is a complex but well-organized multi-step process. In the first reaction, called licensing or pre-replicative complex (pre-RC) assembly, the core of the replicative helicase, the Mcm2-7 complex, is loaded onto replication origins as an inactive double hexamer ((Coster & Diffley, 2017; Evrin et al., 2009; Miller, Locke, Greiwe, Diffley, & Costa, 2019; Remus et al., 2009; Ticau et al., 2017; Ticau, Friedman, Ivica, Gelles, & Bell, 2015)). Next, two essential protein kinases, Dbf4-dependent kinase (DDK), CDK-dependent kinase (CDK), and Mcm10, promote activation of the replicative helicase ((Deegan, Yeeles, & Diffley, 2016; Douglas, Ali, Costa, & Diffley, 2018; Heller et al., 2011; Kanke, Kodama, Takahashi, Nakagawa, & Masukata, 2012; Looke, Maloney, & Bell, 2017; Miyazawa-Onami, Araki, & Tanaka, 2017; Muramatsu, Hirai, Tak, Kamimura, & Araki, 2010; Tanaka & Araki, 2013; Tanaka, Nakato, Katou, Shirahige, & Araki, 2011; Tanaka et al., 2007; van Deursen, Sengupta, De Piccoli, Sanchez-Diaz, & Labib, 2012; Watase, Takisawa, & Kanemaki, 2012; Zegerman & Diffley, 2007)). In this reaction, called firing, the inactive Mcm2-7 double hexamer is converted into an active helicase, the CMG (Cdc45–Mcm2–7–GINS) complex. Finally, the active replication fork is assembled, and nascent DNA is synthesized (elongation).

Chromatin structure is proposed to influence these steps of DNA replication. For example, in mammalian cells, chromatin regulatory factors such as Hbo1, Snf2, GRWD1, and PR-Set7 play roles in promoting licensing (Aizawa, Sugimoto, Watanabe, Yoshida, & Fujita, 2016; Brustel et al., 2017; Chadha & Blow, 2010; Sugimoto et al., 2015; Sugimoto, Yugawa, Iizuka, Kiyono, & Fujita, 2011). They may act by helping to exclude nucleosomes from future Mcm2-7 loading sites, as chromatin-loaded Mcm2-7 double hexamers and nucleosomes occupy their positions on chromatin in a mutually exclusive manner (Berbenetz, Nislow, & Brown, 2010; Eaton, Galani, Kang, Bell, & MacAlpine, 2010). During the elongation reaction, nucleosomes in front of the replication fork must be removed, and then they re-assemble after fork passage. For example, FACT (facilitates chromatin transcription), INO8IEW1a, Gcn5 and Esa1 are known to play an important role in the elongation step both *in vivo* and *in vitro* (Devbhandari, Jiang, Kumar, Whitehouse, & Remus, 2017; Foltman et al., 2013; Kurat, Yeeles, Patel, Early, & Diffley, 2017; Tan, Chien, Hirose, & Lee, 2006). By contrast, the role of chromatin regulatory factors in the firing reaction remains elusive.

Because eukaryotic cells have multiple replication origins, each with their own timing for activation, the firing reaction occurs throughout the S-phase of the cell cycle (reviewed in (Rhind & Gilbert, 2013)), i.e., some origins fire early in S-phase, some fire late, and some do not fire if the replication fork passes through them before firing occurs; the last category of origins are considered ‘dormant’. In a previous study, we showed that Sld3, Sld7, and Cdc45 are loaded onto the pre-RC in a DDK-dependent manner as the first step of firing. Because these factors are much less abundant than replication origins, they are limiting for firing; accordingly, in the budding yeast *Saccharomyces cerevisiae*, simultaneous overexpression of Sld3–Sld7–Cdc45 (3-7-45) affects the firing pattern of origin activation (Tanaka et al., 2011). We observed that untimely activation of dormant origins in the silenced locus *HMLα* occurred under overexpression of 3-7-45. Here, we show that 3-7-45 overexpression also affects the silencing of *HMLα*. These effects depend on the interaction of Sld3 and Esa1, the catalytic subunit of the histone acetyltransferase complex NuA4, which is also called Kat5 (K-acetyl transferase 5). Our data reveal a novel level of firing regulation mediated by the interaction between the firing factor Sld3 and the chromatin regulator Esa1.

## RESULTS

### Simultaneous expression of Sld3–Sld7–Cdc45 counteracts HMLα silencing

In a previous analysis, we showed that the simultaneous overexpression of Sld3, Sld7, and Cdc45 (3-7-45 expression) promotes early firing of many origins, including late and dormant origins (Tanaka et al., 2011). We also observed activation of dormant origins located in and near the silenced locus *HMLα*, such as *ARS301, ARS302, ARS303*, and *ARS320* (Vujcic, Miller, & Kowalski, 1999) (Fig. S1A and B). Therefore, we asked whether the expression of 3-7-45 also affects the silencing status of *HMLα*. To address this question, we used a strain in which wild-type *URA3* is inserted into *HMLα* (*hmlα::URA3*) (Singer & Gottschling, 1994) (Fig. 1A and B). Due to this *URA3* insertion, the silencing status of *HMLα* can be easily monitored: when *HMLα* is silenced, cells are Ura^-^ and 5-FOA–resistant (Fig. 1C, top row: *hmlα::URA3);* conversely, when silencing is lost, as in a *sir* mutant, cells are Ura^+^ and 5-FOA-sensitive (Fig. 1C, bottom two rows: *Δsir1* and *Δsir2*). When 3-7-45 was overexpressed, cells exhibited an intermediate phenotype: they grew better than vector control on SC-Ura, and growth was weakly inhibited on 5-FOA (Fig. 1D, third and fourth rows: shown as ‘+ *GALp-3-7-45*’). Interestingly, the strength of the ‘anti-silencing’ effect caused by the 3-7-45 expression increased with the integrated copy number of the GALp-3-7-45 plasmid (Fig. 1C, third and fourth rows: the numbers of integrated copies of the plasmid are shown in parentheses as ‘x1’ and ‘x2’, respectively), suggesting that 3-7-45 is directly involved in this phenomenon. Together, these observations indicate that the expression of 3-7-45 exerts an anti-silencing effect, and that the strength of this effect is related to the expression levels of 3-7-45.

**Figure 1.**
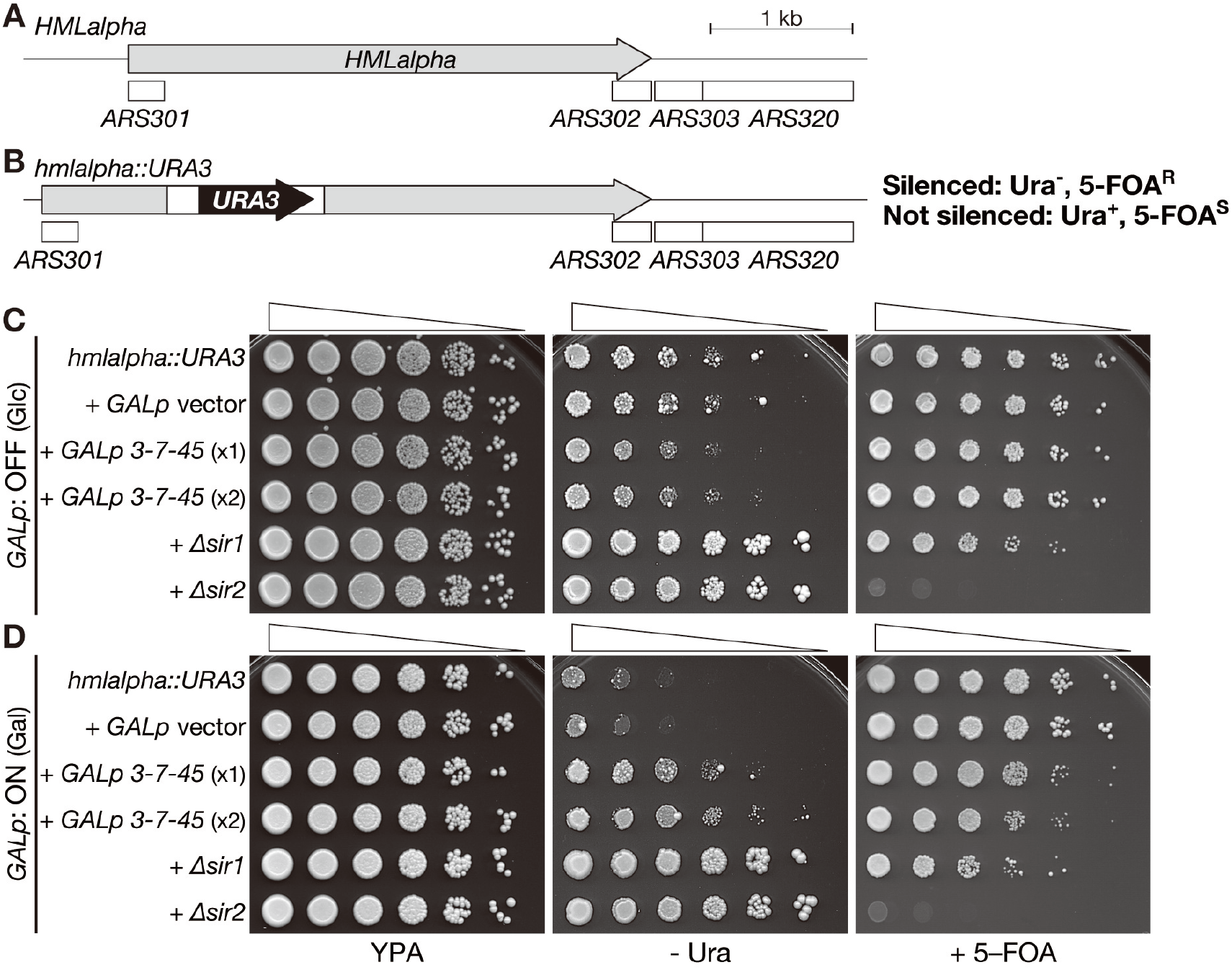
Expression of 3-7-45 induces an anti-silencing effect at *HMLα*. **A.** Schematic of the *HMLα, ARS301, ARS302, ARS303*, and *ARS320*. Positions of each element are based on those in the Saccharomyces Genome Database, SGD (https://www.yeastgenome.org). **B.** Schematic of the *HMLα::URA3* construct. **C.** *HMLα::URA3* cells and *HMLα::URA3* cells harboring the *GALp* vector, 1 or 2 copies of *GALp-3-7-45* (shown as x1 and x2, respectively), and *Δsir1* and *Δsir2* cells were serially diluted, spotted on the indicated plates, and grown at 30°C.

### Anti-silencing effects depend on the replication function of 3-7-45

The Sld3–Sld7–Cdc45 complex functions in origin firing; specifically, it associates with the Mcm2-7 complex at the licensed origin in a DDK-dependent manner during the firing reaction (Heller et al., 2011; Tanaka et al., 2011). In a previous study, we observed that untimely activation of origins caused by 3-7-45 expression required the expression of all three genes; expression of any one or two of them did not induce untimely origin firing except in the case of simultaneous expression of Sld3 and Sld7, which triggered very weak activation of origins (Tanaka et al., 2011). The anti-silencing effect caused by 3-7-45 expression recapitulated this, i.e., the anti-silencing effect was observed only when all three genes were expressed; a very weak anti-silencing effect was observed when Sld3 and Sld7 were simultaneously expressed (Fig. S1C).

If the anti-silencing effect caused by 3-7-45 expression depends on its replication-related function, which involves the association of 3-7-45 with the DDK-phosphorylated Mcm2-7 complex at licensed origins, then the anti-silencing effect must depend on origin licensing. Replication origins are licensed from the end of M-phase until they are activated in S-phase. Therefore, if the duration of time when cells harbor licensed origins is changed, it would affect the anti-silencing effect of 3-7-45. To test this possibility, we used the *Δswe1* and *Δmih1* mutants. Because Swe1 and Mih1 regulate the timing of onset of M-phase via activation of M-phase CDK in opposite manners, and M-phase entry is accelerated in *Δswe1* and delayed in *Δmih1*, the length of G1 phase is longer in *Δswe1* than in the wild type, and shorter in *Δmih1* (Booher, Deshaies, & Kirschner, 1993; Russell, Moreno, & Reed, 1989). The anti-silencing effect of 3-7-45 was strengthened in *Δswe1* cells and weakened in *Δmih1* cells (Fig. 1D and E). On the basis of these observations, we concluded that the anti-silencing effect is a result of the origin association of 3-7-45.

### Sld3 interacts with Esa1

The chromatin regulatory functions of 3-7-45 remain unknown. To begin to investigate this issue, we postulated that any of Sld3, Sld7 and Cdc45 interacts with chromatin regulatory factors. To identify such interactors, we performed systematic yeast two-hybrid (Y2H) assays and found that Sld3 interacts with Esa1, the budding yeast ortholog of lysine acetyl transferase 5 (Kat5), the catalytic subunit of NuA4 histone acetyl transferase (HAT) (Fig. 2A and S2A)(Clarke, Lowell, Jacobson, & Pillus, 1999). Esa1 also exhibited a weak interaction with Sld7 but did not interact with Cdc45 in this assay.

**Figure 2.**
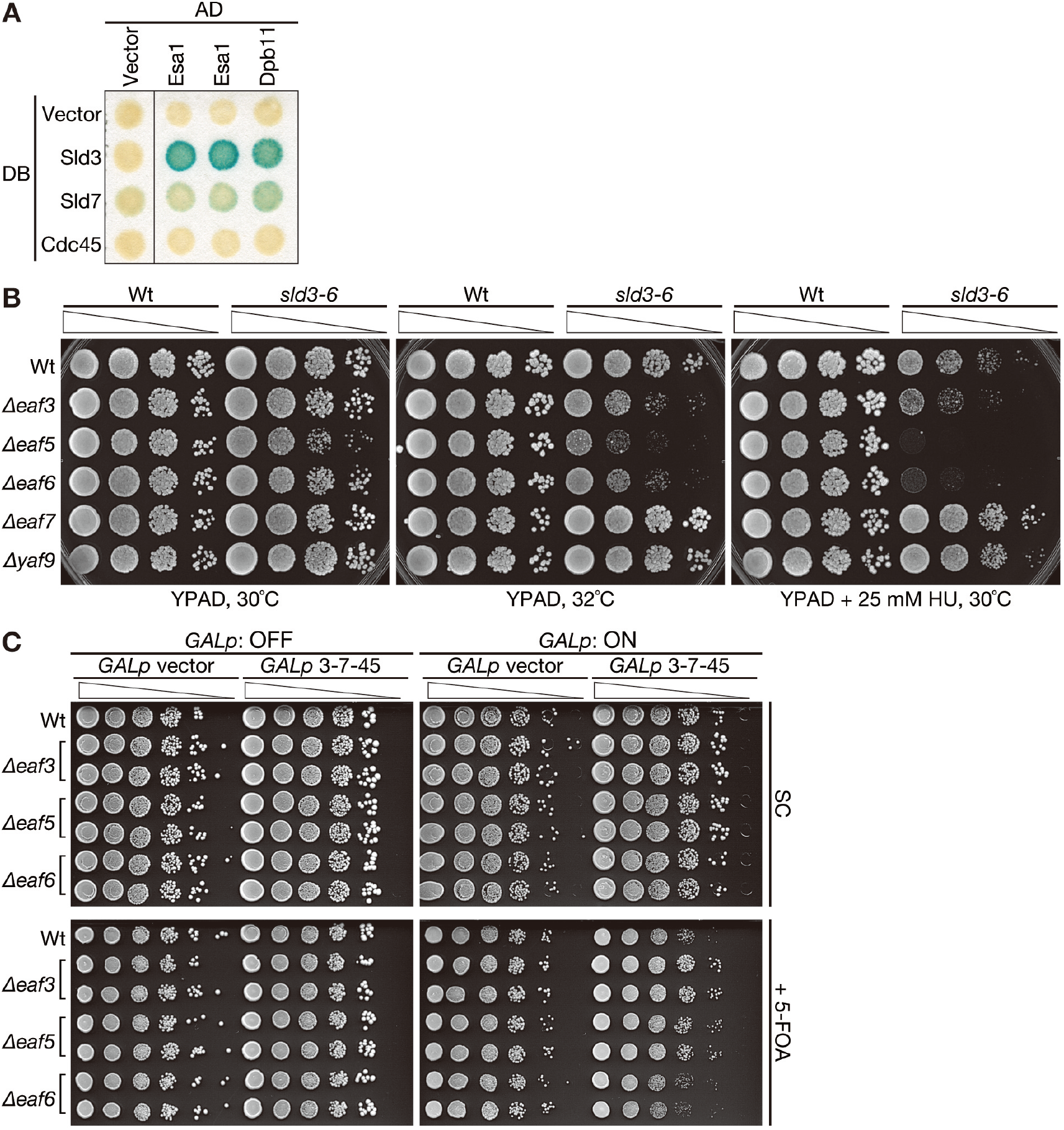
Sld3 interacts with Esa1. **A.** Y2H analysis was performed with plasmids encoding the indicated proteins. AD: activation domain. DB: DNA-binding domain. **B.** *Δeaf3, Δeaf5, Δeaf6, Δeaf7*, and *Δyaf9* cells in the wild type (Wt) or *sld3-6* background were spotted and grown under the indicated conditions. **C.** Wild-type (Wt), *Δeaf3, Δeaf5*, and *Δeaf6* cells harboring the *GALp* vector or *GALp-3-7-45* were spotted and grown under the indicated conditions.

### NuA4 subunits interact genetically with Sld3

NuA4 HAT consists of 13 subunits including Esa1 (Allard et al., 1999). Although Esa1 is essential for cell growth, some subunits of NuA4 are not. To elucidate the functional relationship between Sld3 and NuA4, we deleted several of the non-essential subunits of NuA4, *Δeaf3, Δeaf5, Δeaf6, Δeaf7*, and *Δyaf9* in cells harboring temperature-sensitive alleles of *SLD3, sld3-5*, and *sld3-6* (Kamimura, Tak, Sugino, & Araki, 2001). Some of the mutant alleles exhibited a genetic interaction (Fig. 2B, S2B, and S2C). *Δeaf3, Δeaf5*, and *Δeaf6* exacerbated the sensitivity of the *sld3-6* mutant to high temperature and HU (Fig. 2B). By contrast, *Δeaf7* and *Δyaf9* exhibited an opposite effect in the same background (Fig. 2B and S2B). No such effect was observed in another *sld3-ts* allele, *sld3-5* (Fig. S2C). All of the deletion mutants weakly exacerbated the phenotype of *sld3-5*. This difference between *sld3-5* and *sld3-6* backgrounds might be due to differences in the properties of alleles. Collectively, the results of this experiment revealed a genetic interaction between NuA4 and SLD3.

Based on the existence of a genetic interaction between Sld3 and NuA4 subunits, we next asked whether *Δeaf3, Δeaf5*, and *Δeaf6* affect the anti-silencing effect of 3-7-45, as these three deletions exhibited similar synthetic effects in both *sld3-5* and *sld3-6* backgrounds. Although *Δeaf6* had no obvious effects, *Δeaf3* and *Δeaf5* weakened the anti-silencing effect (Fig. 2D). This indicates that NuA4 somehow contributes to the anti-silencing caused by 3-7-45 expression.

### Deletion of the NuA4 subunit Eaf5 affects the activity of origins in the silenced locus *HMLα*

To further elucidate the genetic interactions between Sld3 and NuA4 subunits, we asked whether NuA4 has an effect on the activities of dormant origins located in the silenced locus *HMLα*. To answer this question, we performed a plasmid loss assay with centromeric plasmids harboring the early replication origin *ARS1*, a short fragment derived from *HMLα* that only contains *ARS301*, or the whole region of *HMLα* plus *ARS302-320* (hereafter denoted as *ARS1, ARS301*, and *ARS301-320*’, for details, see Fig. S3A and B). The *ARS301*-containing short fragment has characteristics of a late replication origin even in a plasmid context (Bousset & Diffley, 1998; Santocanale & Diffley, 1996), probably due to its chromatin structure. Therefore, we first investigated whether the *HMLα*-derived fragments were silenced on the plasmid. The *URA3* reporter gene, which was inserted proximal to or between the *HMLα*-derived origins, was silenced in the *ARS301* and *ARS301-320* plasmids, but not in the *ARS1* plasmid (Fig. S3C). We concluded that *HMLα*-derived fragments are silenced in the plasmid context, as in the chromosome context.

*ARS301* and *ARS301-320* plasmids were lost at higher rates than the *ARS1* plasmid (Fig. 3A), suggesting poorer DNA replication of these origins. Importantly, this high plasmid loss rate was suppressed by deletion of *SIR2*, which encodes silencing factor (Rine & Herskowitz, 1987), *Δsir2* (Fig. 3A), implying that the higher loss rate of *ARS301* and *ARS301-320* plasmids was related to silencing. Deletion of the NuA4 subunit Eaf5 increased the loss rate of *ARS301* and *ARS301-320* plasmids, but not that of the *ARS1* plasmid (Fig. 3A), indicating that the deletion interfered with replication of these plasmids. Conversely, 3-7-45 expression suppressed the high loss rate of *ARS301* and *ARS301-320* plasmids (Fig. 3B). Finally, the combination of *Δeaf5* and 3-7-45 expression resulted in an intermediate loss rate for the *ARS301 and ARS301-320* plasmids (Fig. 3C). Therefore, we concluded that the NuA4 complex, at least Eaf5, positively affects not only the anti-silencing effect of 3-7-45 but also the activation of dormant origins in *HMLα*.

**Figure 3.**
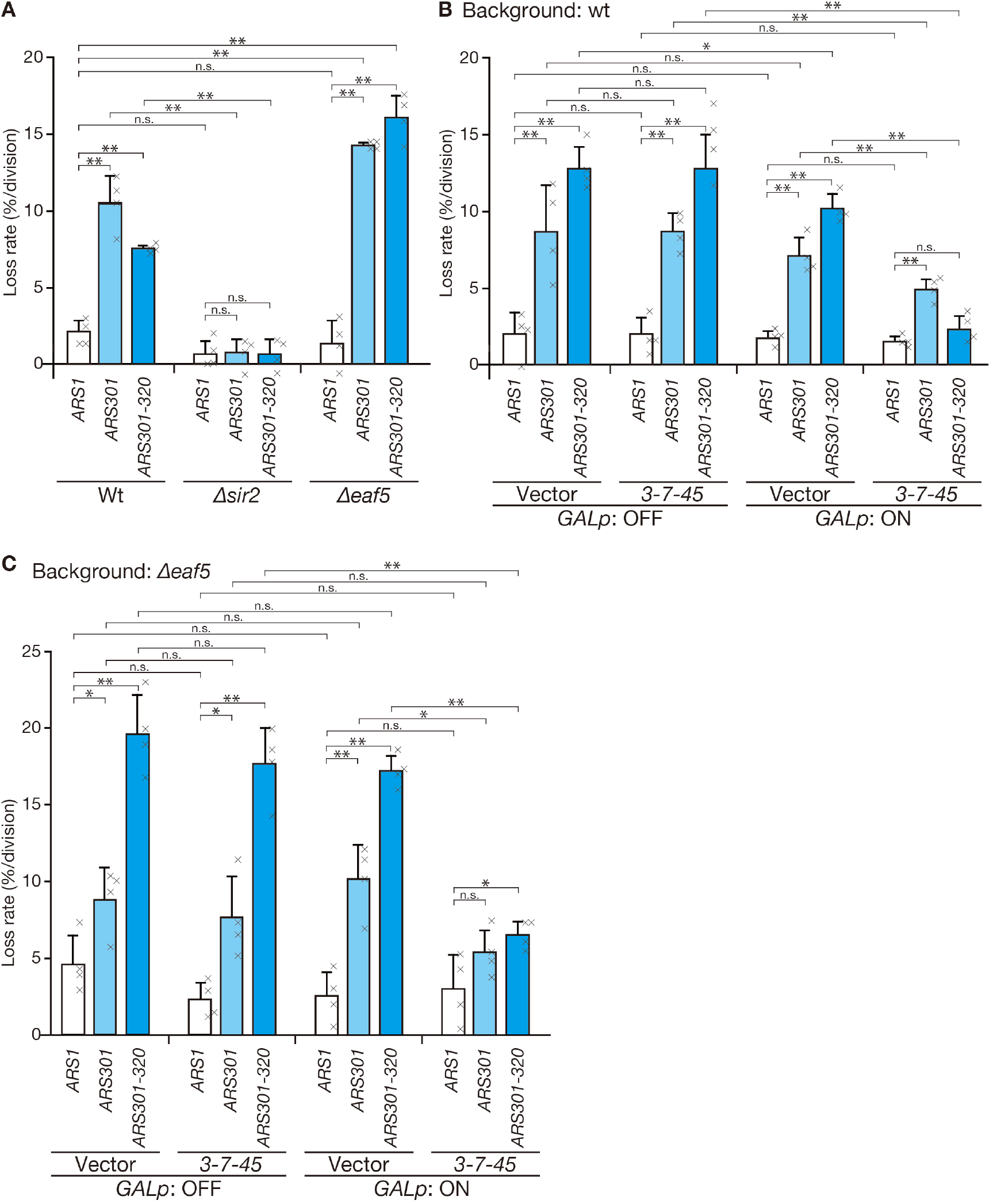
Plasmids harboring *HMLα* origins replicate more efficiently when 3-7-45 is expressed, and less efficiently in the *Δeaf5* background. **A.** Plasmid loss rates were measured using plasmids harboring the indicated ARSs (*ARS1*: pST1969; *ARS301:* pST1970; and *ARS301-320:* pST1971; see Supplementary Fig. S3A and D for details). **B.** Plasmid loss assay was performed with host strains harboring either *GALp* vector or *GALp-3-7-45*. **C.** Plasmid loss assay was performed with *Δeaf5* strains harboring either *GALp* vector or *GALp-3-7-45*. For each experiment, the average of four independent results is shown. Error bars: SD. Statistical significance in the differences was tested by the Student’s t-test. **: P < 0.01, *: 0.01 < P < 0.05, and n.s. (not significant): P > 0.05.

### The Sld3–Esa1 interaction is important for the anti-silencing effects of 3-7-45 and the activation of replication origins in the silenced locus *HMLα*

Experiments with *Δeaf5* suggested a positive role for NuA4 in the anti-silencing effect of 3-7-45 and the activation of dormant origins in the silenced locus *HMLα*. However, deletions of non-essential subunits revealed a complex genetic interaction with *sld3-6* (Fig. 2B and S2B), and no interaction between Eaf5 and Sld3 could be detected in the Y2H assay (Fig. S4A). Because an Esa1–Sld3 interaction was identified in the same assay (Fig. 2 and S4A), we investigated the role of that interaction, rather than the NuA4 complex itself, in the role of 3-7-45 in anti-silencing and DNA replication of dormant origins in the silenced locus. For this purpose, we isolated Sld3 mutants with diminished interaction with Esa1. Because Sld3 is essential for growth, we sought to isolate viable mutants for further analysis. Ultimately, we obtained three such mutants, *sld3^Esa1(-)^−1, −2*, and *-3* (Material and Methods, Fig. 4A). These three Sld3^Esa1(-)^ mutants exhibited a significant reduction in the Esa1 interaction in the Y2H assay; in particular, Sld3^Esa1(-)^-3 seemed to lose the interaction completely. Importantly, all mutants maintained interactions with other replication factors such as, Sld7, Cdc45, Dpb11, Orc2, and Mcm6 at levels similar to those in the wild type (Fig. 4A), indicating that they only diminished or lost the ability to interact with Esa1.

**Figure 4.**
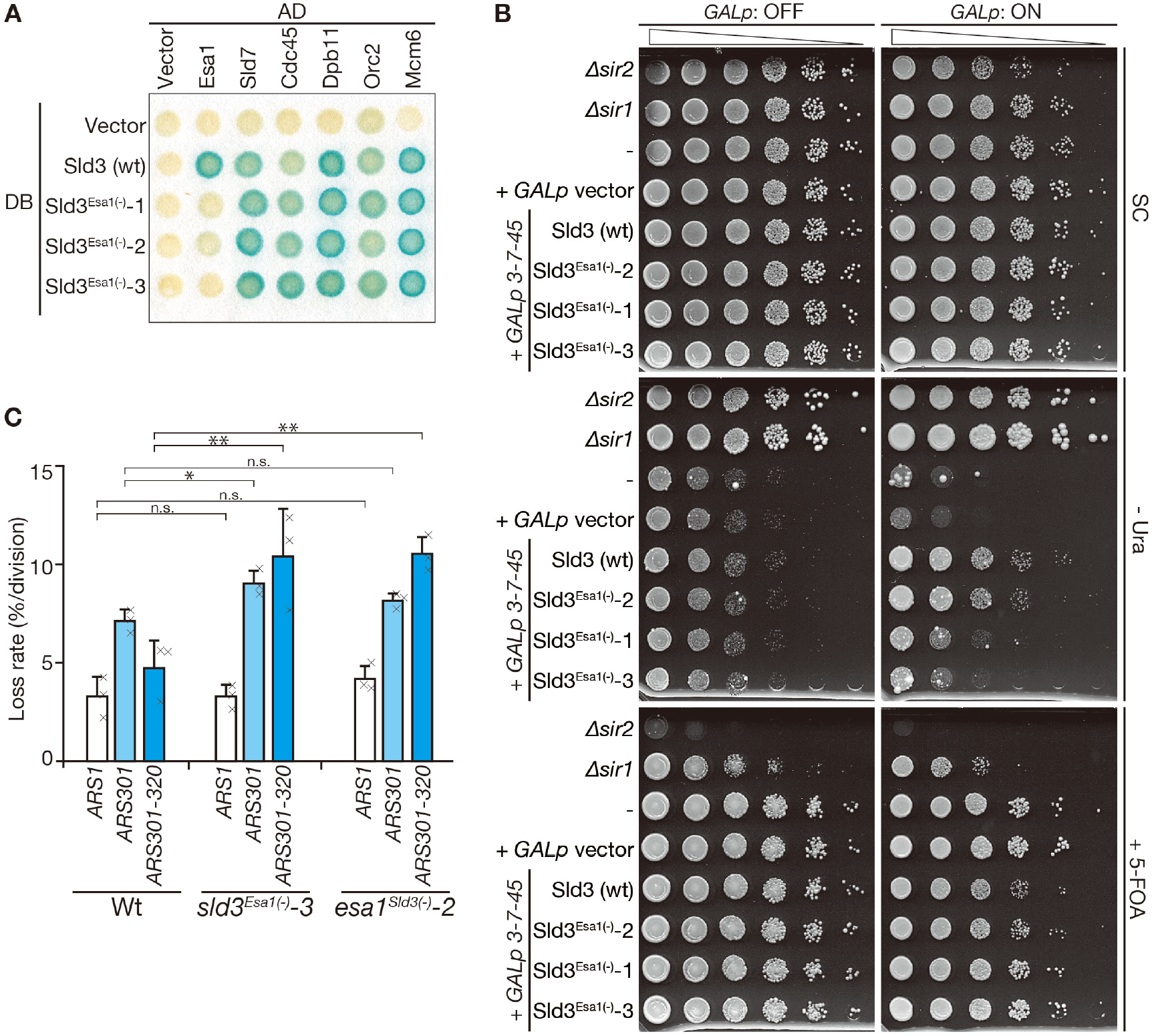
Mutants that diminish the Sld3–Esa1 interaction exhibit weaker anti-silencing effects and higher plasmid loss of *HMLα* origins. **A.** Isolation of Sld3 mutants with diminished interactions with Esa1. Y2H analysis was performed with the plasmids encoding the indicated proteins. AD: activation domain. DB: DNA-binding domain. **B.** Cells expressing Sld3 wild-type or Sld3^Esa1(-)^ mutant proteins with Sld7 and Cdc45 were serially diluted, spotted, and grown on the indicated plates. **C.** Plasmid loss assay was performed in the *sld3^Esa1(-)^-3* and *esa1^Sld3(-)^-3* strains. The average of three independent results is shown. Error bars: SD. Statistical significance in the differences was tested by the Student’s t-test. **: P < 0.01, *: 0.01 < P < 0.05, and n.s. (not significant): P > 0.05.

Mapping of mutated amino acids in Sld3^Esa1(-)^ mutants revealed that all three mutants have amino acid substitutions in the Sld3/Treslin homology domain, which is located in the middle of the Sld3 protein and conserved in Sld3 orthologs (Sanchez-Pulido, Diffley, & Ponting, 2010)(Fig. S4B). Moreover, both Sld3^Esa1(-)^-1 and −3 harbor an extra amino acid substitution in the C-terminal region, but these mutations seemed not to affect the Sld3^Esa1(-)^ phenotype, as mutations in the Sld3/Treslin homology domain (N178F and N189Y in the Sld3^Esa1(-)^-1 and P157T, T193S, and V264L in the Sld3^Esa1(-)^-3) were sufficient for a diminished interaction with Esa1, and a single amino acid substitution in the C-terminal region alone (S459L in the Sld3^Esa1(-)^-1 and P623Q in the Sld3^Esa1(-)^-3) did not affect the Sld3–Esa1 interaction (data not shown). These findings imply that the Sld3/Treslin homology domain is important for the Sld3–Esa1 interaction.

To evaluate the importance of the Sld3–Esa1 interaction, we expressed Sld3^Esa1^(-) simultaneously with Sld7 and Cdc45; we observed that the anti-silencing effect was weaker in cells expressing Sld3^Esa1(-)^ than in those expressing wild-type Sld3 (Fig. 4B). This effect was roughly proportional to the decrease in the Y2H signal, as this effect was stronger with the Sld3^Esa1(-)^-1 and −3 than that with Sld3^Esa1(-)^-2, which still shows very weak interaction with Esa1 in the Y2H assay (Fig. 4A and 4B). These observations indicate that the Sld3–Esa1 interaction is important for the anti-silencing effect caused by 3-7-45 expression. To observe the effect on the activity of dormant origins in *HMLα*, we performed a plasmid loss assay. The loss rate of both the *ARS301* and *ARS301-320* plasmids was higher in the *sld3^Esa1(-)^-3* background than in the wild type, although the loss rate of the *ARS1* plasmid was not changed.

We also isolated two viable Esa1 mutants with diminished interactions with Sld3, *esa1^Sld3(-)^-1* and *-2* (Fig. S4C). Both mutants had multiple amino acid substitutions in the middle of the Esa1 protein, and at least one substitution was located in the MYST-type HAT domain. Of the 445 amino acids in Esa1, the N-terminal portion (aa 1–146) was dispensable for the Sld3 interaction, and most of the remaining part of the protein (aa 147–445) was a MYST-type HAT domain (aa 162–433) (Fig. S4A). Therefore, the MYST-type HAT domain is important for the interaction with Sld3. As in the case of *sld3^Esa1(-)^-3*, the loss rate of *ARS301* and *ARS301-320* plasmids was elevated in *esa1^Sld3(-)^-2* (Fig. 4C). Collectively, these data indicate that the Sld3–Esa1 interaction has a positive effect on the activation of origins in the silenced locus.

### The Sld3–Esa1 interaction is important in the activation of not only dormant origins, but also late origins, on the chromosome

As described above, 3-7-45 expression can promote the untimely activation of dormant origins in *HMLα* in a chromosomal context. Therefore, we further evaluated our *sld3^Esa1(-)^* and *esa1^Sld3(-)^* mutants in the same situation. When Sld3^Esa1(-)^-7-45 was expressed, although the activation of early origins such as *ARS305* and *ARS306* was not affected, the untimely activation of dormant origins in the *HMLα* locus, *ARS301* and *ARS302-320*, was diminished (Fig. 5A). Surprisingly, in the both mutants, untimely activation of late replication origins *ARS501* and *ARS1410* were also diminished (Fig. 5A). Similar results were also observed when 3-7-45 was expressed in the *esa1^Sld3(-)^* mutants, although the effect was not remarkable in the *esa1^Sld3(-)^-1* (Fig. 5B). These data suggest that the Esa1–Sld3 interaction plays a role not only in the activation of dormant origins in the silenced locus, but also in the activation of late origins.

**Figure 5.**
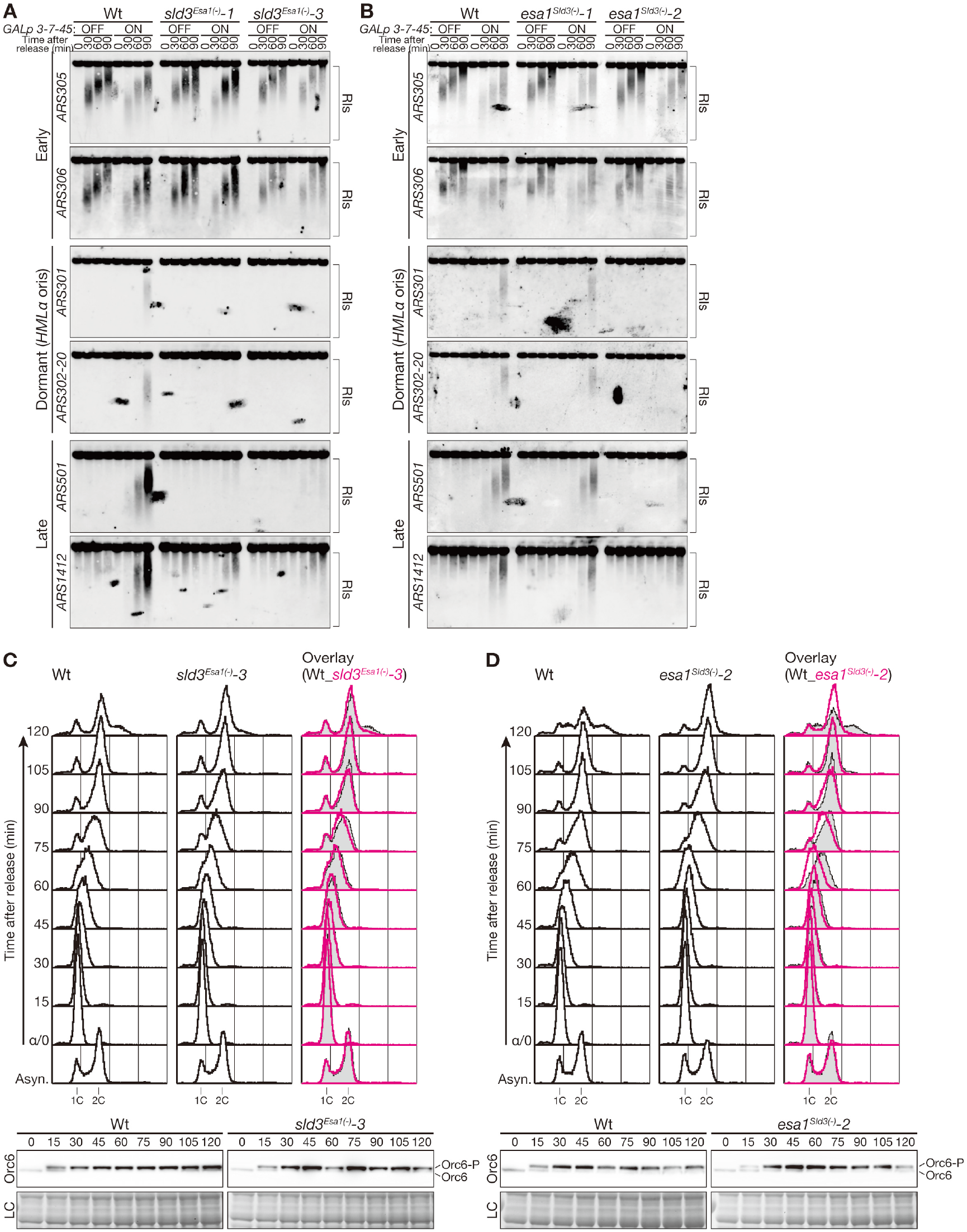
Untimely activation of origins is diminished in mutants that diminish the Sld3–Esa1 interaction. **A.** Wild-type, *sld3^Esa1(-)^-1*, and *sld3^Esa1(-)^-3* cells in which 3-7-45 was expressed were arrested in G1 with α factor, and the culture was split into two. 3-7-45 was induced in one half but not in the other; the cultures were then released into the media containing HU, and samples were collected at the indicated times. DNA was extracted from the samples and analyzed using alkaline gels. Replication intermediates (RIs) were detected using ARS-specific probes. Please note that in *sld3^Esa1(-)^* cells, the corresponding Sld3^Esa1(-)^ protein was expressed rather than wild-type Sld3. Profiles of DNA contents of samples are shown in Supplementary Fig. S5. **B.** RIs from various ARSs in wild-type, *esa1^Sld3(-)^-1*, and *esa1^Sld3(-)^-2*cells in which 3-7-45 were expressed were analyzed as in A. **C.** Wild-type and *sld3^Esa1(-)^-3* cells were arrested in G1 with α factor and released in fresh YPAD medium containing 25 mM HU. Samples were collected every 15 min after release and analyzed by flow cytometry (top) and western blotting (bottom): Orc6 proteins were detected with SB49 monoclonal antibody (Liang & Stillman, 1997); LC (loading control) shows the corresponding region of the membrane stained with Ponceau-S. **D.** Wild-type and *esa1^Sld3(-)^-2* cells were analyzed as in C.

Finally, we asked whether the Esa1–Sld3 interaction affects DNA replication under the normal condition by monitoring DNA replication in cells harboring the *sld3^Esa1(-)^-3* or *esa1^Sld3(-)^-2* allele as the sole copy of the respective genes. Although overall progression of S-phase was largely unaffected when cells were synchronously released from G1 arrest, a very small delay might occur in late S-phase (Fig. S5C and D). Therefore, to further clarify this point, we monitored DNA replication of the *sld3^Esa1(-)^-3* and *esa1^Sld3(-)^-2* cells in the medium containing a low-dose of hydroxyurea (HU). Because high-dose of HU (200 mM) blocks DNA replication by preventing the dNTP pool expansion (Koc, Wheeler, Mathews, & Merrill, 2004) (for example, see Fig. S5A and S5B), low-dose of HU (25 mM) was employed. Under the such mild condition, S-phase progression is allowed but slows down (Fig. 5C and 5D). Judging from the phosphorylation status of Orc6, a well-known substrate of S- and M-phase CDK (Liang & Stillman, 1997; Tanaka et al., 2011), S-phase CDK is activated from 15 min after the release from G1 arrest and DNA replication seemed to initiate at the same time in all strains (Fig. 5C and 5D). However, the progression of S phase in later time points (45-105 min) in *sld3^Esa1(-)^-3* and *esa1^Sld3(-)^-2* cells were slower than that of wild-type cells (Fig. 5C and 5D). These suggest that the firing of early origins was unaffected, but that of late origins was affected, in *sld3^Esa1(-)^-3* and *esa1^Sld3(-)^-2* cells.

## DISCUSSION

In this study, we showed that 3-7-45 expression affects not only the firing of dormant origins in the silenced locus *HMLα*, but also the silencing status of the locus. These effects depended on the interaction of Sld3 and Esa1. The Sld3–Esa1 interaction also affected the activation of late origins. Therefore, our data reveals a novel mechanism regulating the firing of replication origins.

### The anti-silencing effect caused by 3-7-45 expression is transient

Chromatin silencing is maintained over generations. In this study, by monitoring the activity of a *URA3* gene inserted into the *HMLα* locus, we found that 3-7-45 expression counteracted the silencing of the locus. This effect was strengthened in the *Δswe1* mutant, which has a longer G1 phase than the wild type, and the opposite effect was observed in the *Δmih1* mutant, which has a shorter G1 phase than the wild type. 3-7-45 transiently associates with the Mcm2-7 complex on licensed origins when they fire, and origins are licensed in the G1 phase, implying that the anti-silencing effect of 3-7-45 is related to their association to licensed replication origins.

On 5-FOA plates, colonies expressing 3-7-45 were smaller than those expressing vector control, indicating that the level of Ura3 was higher in the 3-7-45 cells; by contrast, the number of colonies formed was largely unaffected (Fig. 1D). Therefore, the anti-silencing effect of 3-7-45 seems to be transient, rather than representing a switch in status. When Sld3^Esa1(-)^ mutants were expressed, colony size increased, but once again the number of colonies was not affected (Fig. 4B, +5-FOA panel). Therefore, we propose the following model: when 3-7-45 is associated with licensed origins, local chromatin structure is transiently changed via the function of Esa1. Although our data cannot strictly exclude the possibility that Esa1 locates silenced origins prior to the association of Sld3, it would be difficult to explain the transient effect of 3-7-45 expression in such a situation.

Interestingly, in the *in vitro* replication assay with chromatinized template, Esa1 accumulated on template chromatin DNA when S-phase extract was added. This requires Mcm2-7 loading and subsequent DDK treatment (Kurat et al., 2017). Because Sld3 associates with the Mcm2-7 complex at licensed origins in a DDK-dependent manner (Deegan et al., 2016; Heller et al., 2011; Tanaka et al., 2011), these *in vitro* data are consistent with our model.

In chromatin immunoprecipitation (ChIP) assays, we tried to detect association with Esa1 or a reduction in the level of the silencer protein Sir2 at the *HMLα* locus following 3-7-45 expression, but we detected no changes (data not shown). Because Sld3 transiently associates with licensed origins during the firing reaction, the association of Esa1 with silenced origins might occur in a very short window, and ChIP analysis might not be sensitive enough to detect such transient changes.

Genome silencing is stably maintained throughout the cell cycle and over generations. Hence, our finding that temporary anti-silencing, or fluctuation of silencing, can occur when an origin fires reveals a novel mode of silencing regulation.

### Target and the role of Esa1 in the firing reaction

Esa1, a lysine acetyltransferase and the catalytic subunit of the NuA4 complex, acetylates four conserved lysines in the histone H4 N-terminal tail at positions 5, 8, 12, and 16 (Dhar, Gursoy-Yuzugullu, Parasuram, & Price, 2017). Therefore, transient recruitment of Esa1 by Sld3 might transiently change the modification levels of local proteins. Using ChIP with the specific antibodies against H4K5Ac, H4K8Ac, H4K12Ac, and H4K16Ac raised by the Kimura group (Hayashi-Takanaka et al., 2015), we tried to detect changes in these modifications in 3-7-45 expressing cells. However, as in the case of the Esa1-ChIP, we could not detect significant changes, probably because of technical limitations. Further analysis that is aimed at identifying Esa1 targets, including non-histones, will be necessary to understand the precise role of Esa1 in the firing reaction. The aforementioned *in vitro* replication assay (Kurat et al., 2017) would be very useful for elucidating the function of Esa1 in the firing reaction.

### Is the full NuA4 complex required for anti-silencing and untimely activation of *HMLα* origins caused by 3-7-45 expression?

Budding yeast NuA4 consists of 13 unique subunits, including Esa1(Allard et al., 1999). We tested the genetic interactions between NuA4 and Sld3 by combining deletions of non-essential subunits of NuA4 (*Δeaf3, Δeaf5, Δeaf6, Δeaf7*, and *Δyaf9*) with *sld3-ts* alleles. Although we detected genetic interactions (Fig. 2 and S2), the strengths and directions of these interactions were complex. Although this might be partly due to differences between *sld3-5* and *sld3-6* (see below), the structure of the NuA4 complex might also affect the phenotype. For example, the 13 subunits of NuA4 can be separated into subcomplexes, one of which is the TINTIN triad consisting of Eaf3, Eaf5, and Eaf7 (Cheng & Cote, 2014; Rossetto et al., 2014). *Δeaf3* and *Δeaf5* exacerbated the growth defects of *sld3-6* to different extents, whereas *Δeaf7* suppressed the growth defects (Fig. 2B and S2B). By contrast, in *sld3-5, Δeaf5* and *Δeaf7* weakly exacerbated the growth defects, whereas *Δeaf3* did not (Fig. S2C). Moreover, although both *Δeaf5* and *Δeaf6* exacerbated the phenotypes of *sld3-5* and *sld3-6*, neither Eaf5 nor Eaf6 interacted with Sld3 in the Y2H assay. Together, these findings suggest the existence of different role(s) for NuA4 subunits or other subcomplexes.

### Conservation of the Esa1–Sld3 interaction and its impact on genome duplication

We isolated Sld3^Esa1(-)^ and Esa1^Sld3(-)^ mutants in which the Sld3–Esa1 interaction was diminished. All mutants had at least one amino acid substitution in their conserved domains: the Sld3/Treslin homology domain in Sld3 and the MYST-type HAT domain in Esa1. The Sld3/Treslin homology domain is a conserved Cdc45-binding domain that contains 12 helices (Itou, Muramatsu, Shirakihara, & Araki, 2014). All Sld3^Esa1(-)^ mutants had an amino acid substitution in the helix 2 (aa 176–195). The helix 2 contains basic amino acids that are conserved and form a basic patch on the surface of the Sld3/Treslin homology domain with basic amino acids in the helix 12. This basic patch does not seem to be the Cdc45 association domain, because the Cdc45 association was unaffected in the KR2E mutation of the Sld3/Treslin homology domain, in which five basic amino acids in the helix 2 and 12 were replaced by glutamic acid (Itou et al., 2014). Therefore, helix 2 in the Sld3/Treslin homology domain might be the conserved Esa1 interaction domain.

We used two temperature-sensitive alleles of *sld3*, *sld3-5* and *sld3-6* (Kamimura et al., 2001), to analyze the interaction between Sld3 and NuA4. A genetic interaction was observed in both mutants; however, the interaction in *sld3-5* was much weaker than that in *sld3-6* (Fig. 2 and S2). The amino acid substitutions in Sld3-5 are G125D and F170S, and the substitution in Sld3-6 is E63G (Kamimura et al., 2001). F170 is located in helix 2 in the Sld3/Treslin homology domain. Therefore, if helix 2 is responsible for the association with Esa1, it could explain the difference in the genetic interactions between NuA4 and *sld3-5* and *sld3-6*. Notably in this regard, the KR2E mutant does not support cell growth (Itou et al., 2014), whereas our Sld3^Esa1(-)^ mutants did. Therefore, further investigations are necessary to understand the exact role of the basic patch in the Sld3/Treslin homology domain in the Sld3–Esa1 interaction.

Although the mutation sites of our Esa1^Sld3(-)^ mutants were dispersed throughout the protein (Fig. S2D), both Esa1^Sld3(-)^ mutants have amino acid substitution(s) in the conserved MYST-type HAT domain. Given that both Sld3^Esa1(-)^ and Esa1 ^Sld3(-)^ mutants had amino acid substitutions in their conserved domains, the Sld3–Esa1 interaction might be conserved through evolution.

In this study, we showed that untimely firing of dormant origins and late origins, which is induced by 3-7-45 expression, was diminished in Sld3^Esa1(-)^ and Esa1 ^Sld3(-)^ (Fig. 5). When DNA replication was monitored in cells harboring the *sld3^Esa1(-)^-3* or *esa1^Sld3(-)^-2* allele as the sole copy of the respective genes, overall progression of S-phase was largely unaffected, although a very small delay might occur in late S-phase (Fig. S5C and D). Similar phenotype was observed with the degron allele of *ESA1, esa1-td* (Early, Drury, & Diffley, 2004). Dormant origins have the potential to fire, but under normal conditions do not fire because replication forks pass through before they can (Bartel & Fields, 1995; Santocanale, Sharma, & Diffley, 1999). Late origins fire under normal conditions; however, if firing is slightly delayed, a replication fork could pass through before firing. We speculate that such a situation is arising in our *sld3^Esa1(-)^* and *esa1^Sld3(-)^* mutants and *esa1-td;* therefore, the overall progression of S-phase in these mutants was not strongly affected in our *sld3^Esa1(-)^* and *esa1^Sld3(-)^* mutants (Fig. 5C and 5D). In the budding yeast *S. cerevisiae*, only a small amount of the genome is silenced; by contrast, in mammals, huge sections of the genome are silenced. Because the Sld3–Esa1 interaction may be conserved, future studies should seek to elucidate the role of Sld3–Esa1 in other eukaryotes.

## MATERIALS and METHODS

### Yeast strains and media

*S. cerevisiae* strains are listed in Supplementary Table 1. Cells were grown in the rich medium YPA (1% yeast extract, 2% Bacto peptone, 40 mg/L adenine) supplemented with 2% sugar (glucose, galactose, or raffinose), or in synthetic medium containing 0.67% yeast nitrogen base without amino acids (#291940, Difco Laboratories, Detroit, MI, USA) and 2% glucose, supplemented with the appropriate amino acids and drugs. Cell-cycle block and release experiments with α-factor were performed as described previously (Miyazawa-Onami et al., 2017; Tanaka et al., 2011). All strains used in this analysis are available from the National Bio Resource Project (NBRP) yeast (http://yeast.nig.ac.jp/yeast/top.xhtml).

### Screening of chromatin regulatory factors interact with Sld3, Sld7 and Cdc45

To identify chromatin regulatory factors that interact with Sld3, Sld7 and Cdc45, we cloned 26 genes that were listed by the keyword ‘histone acetyltransferase’ in the Saccharomyces Genome Database web site (https://www.yeastgenome.org/). Each of them were cloned into the vectors for the yeast two-hybrid (Y2H) assay: DNA-binding domain, DB, vector pST1667 and the activation domain, AD, vector pST548 (Tanaka, 2019), respectively, and their capability of interaction with any of Sld3, Sld7 and Cdc45 was tested by the Y2H color assay (Bartel & Fields, 1995).

### Isolation of mutants with diminished interactions between Sld3 and Esa1

To construct the mutant library, *SLD3* and *ESA1* ORF were PCR-amplified with Taq polymerase (MightyAmp ver. 2, Takara Bio Co., Otsu, Japan) and cloned into the *BamHI* sites of the Y2H DB vector pST1667 and the Y2H AD vector pST548 (Tanaka, 2019), respectively, using the In-Fusion enzyme (Takara Bio). Mutant libraries of DB-Sld3 and AD-Esa1 were introduced into the Y2H host TAT-7 strain harboring AD-Esa1 and DB-Sld3, respectively. After 1 day of incubation, the original plates were replicated, and the incubation was continued. After colony formation, cells were lifted from the replicated plates and subjected to the Y2H color assay (Bartel & Fields, 1995). ‘White’ candidates were picked from the original plate and re-streaked for confirmation of the white phenotype. In parallel, whole-cell extracts were prepared from the re-streak and analyzed by western blotting. Candidates generating full-length or near-full-length protein were stored as secondary candidates. Plasmids were recovered from secondary candidates, and the white phenotype in the Y2H color assay was confirmed (tertiary candidates). From tertiary candidates, clones with endogenous promoters were constructed in YCp-type plasmids, and clones that could support cell growth were selected by plasmid shuffling; the resultant mutants were denoted *sld3^Esa1(-)^* and *esa1^Sld3(-)^*.

### Other methods

Flow cytometry was performed as described (Tanaka & Diffley, 2002b). Replication intermediates were detected using alkaline gels as described (Tanaka et al., 2011). Preparation of yeast whole-cell extracts and western blotting were performed as described (Tanaka et al., 2011). Yeast two-hybrid assay was performed as described (Tanaka et al., 2013). The plasmid loss rate was measured as described (Tanaka & Diffley, 2002a)

## Author contributions

ST is responsible for all aspects of the study.

## SUPPLEMENTAL INFORMATION

Supplemental Information includes five figures and one table.

## ACKNOWLEDGEMENTS

We are deeply grateful to M. Miyazawa-Onami and A. Hosoe for their technical assistance, to H. Kimura for antibodies against acetylated histones and to B. Stillman for the SB49 antibody. We also thank H. Araki, M. Yagura, and Y. Tanaka for help and discussion. This work was supported by grants from the Uehara Memorial Foundation; the Institute for Fermentation, Osaka; the Ministry of Education, Culture, Sports, Science and Technology (MEXT); and the Japan Society for the Promotion of Science (JSPS; Grant No. 25131721).

## CONFLICTS of INTEREST

The author have no conflicts of interest associated with this manuscript.

---

**Supplementary Table S1.**
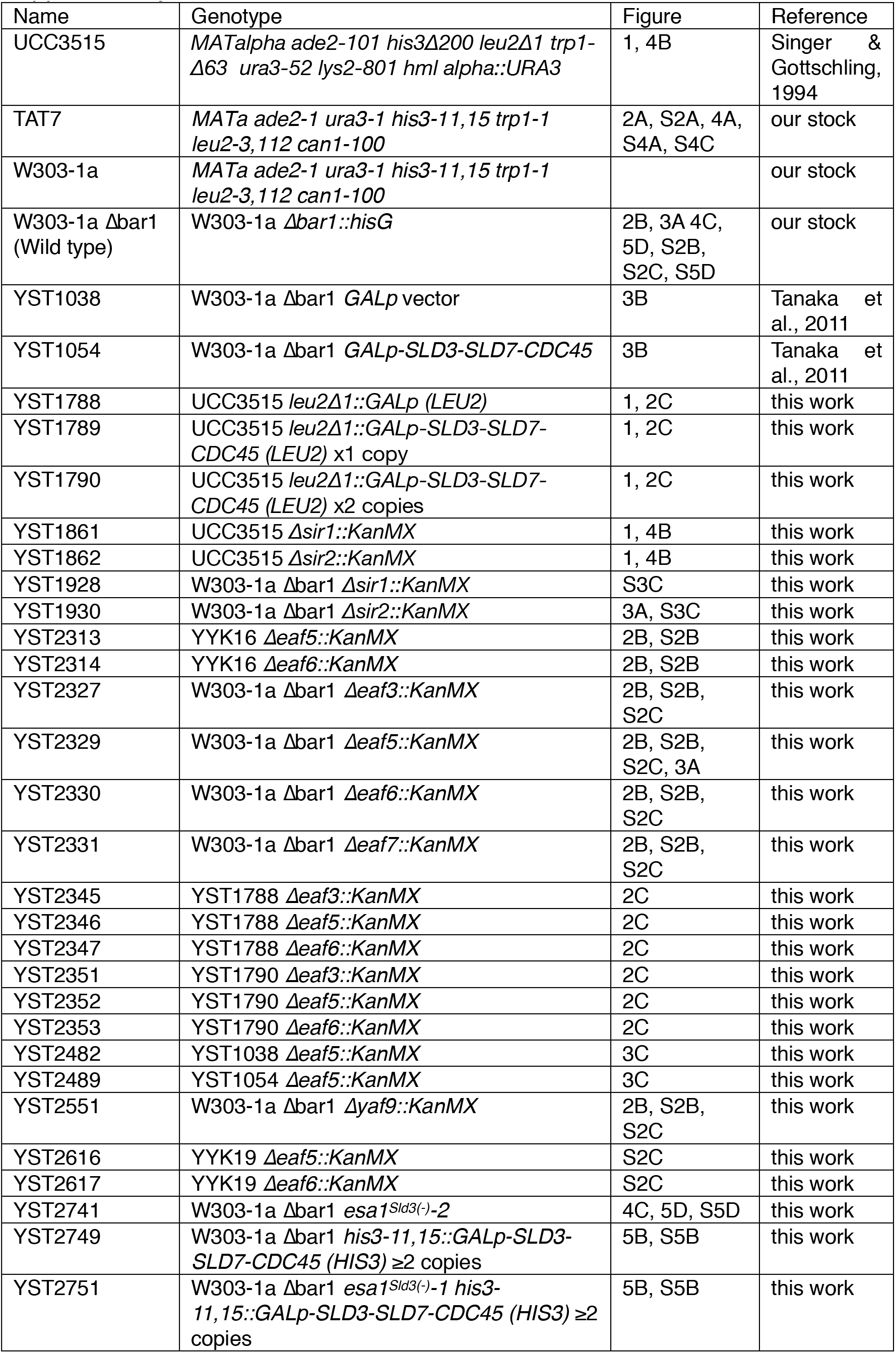
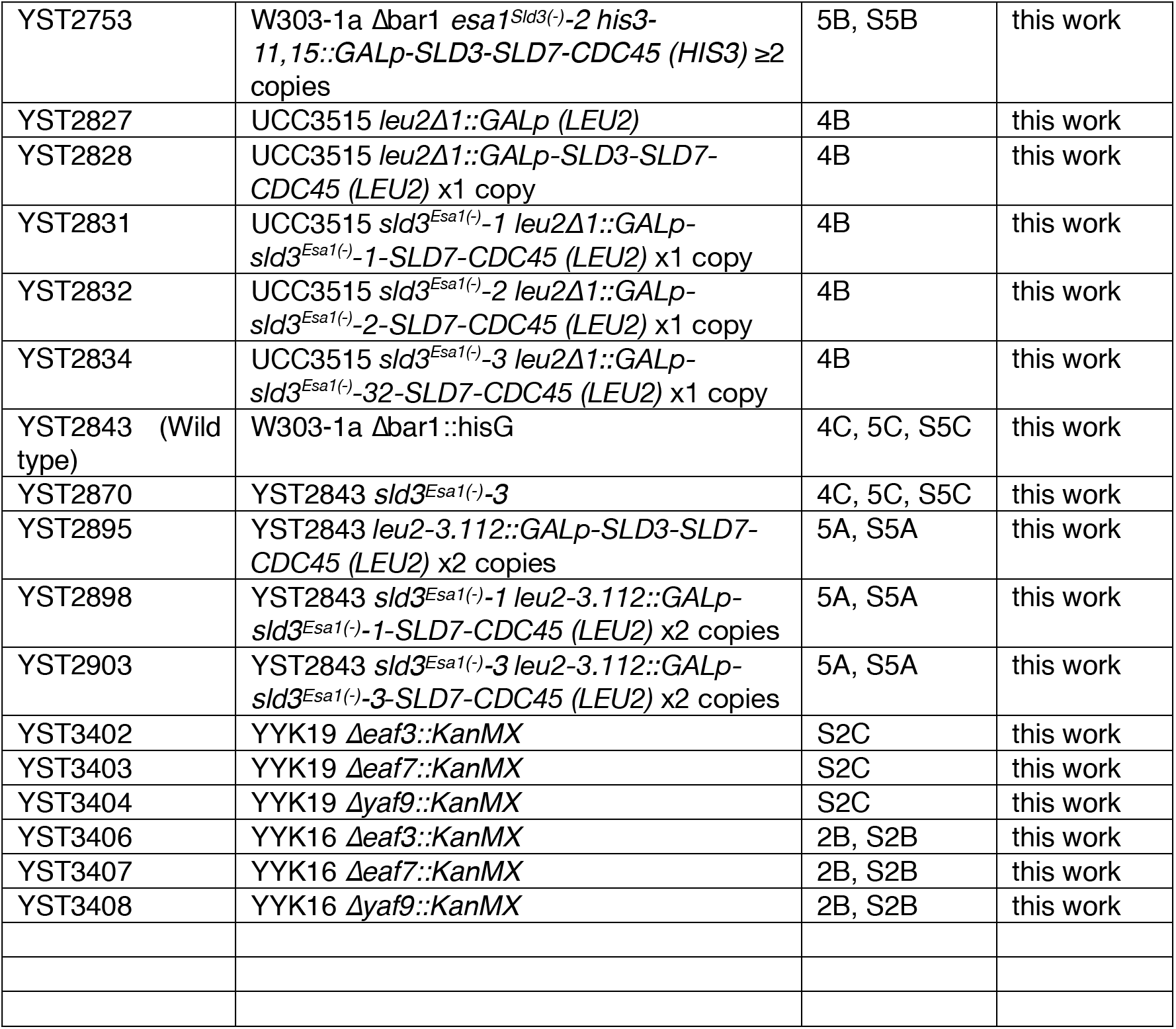
Yeast strains

**Supplementary Figure S1.**
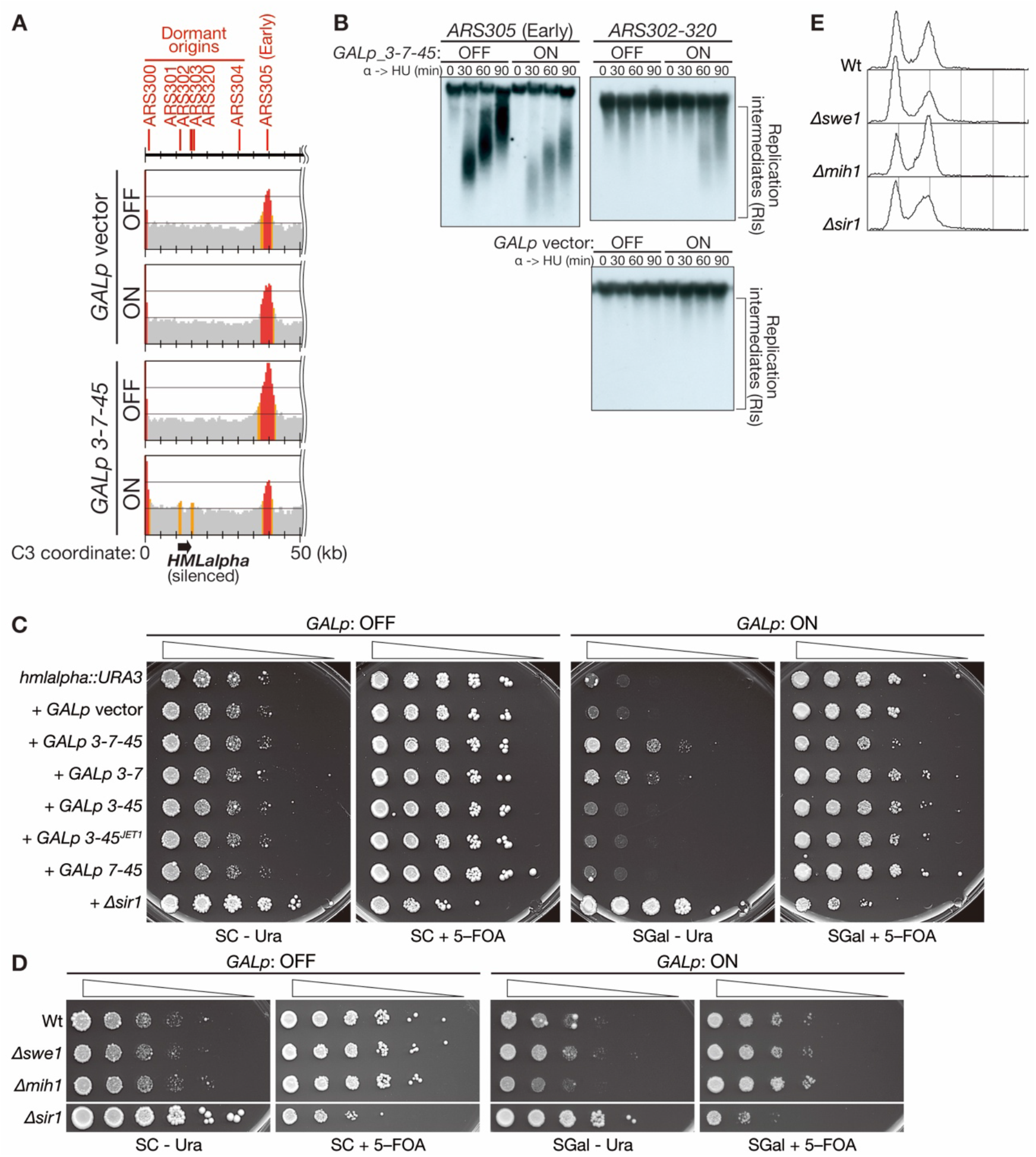
Dormant origins located in and near the silenced locus *HMLα* are activated upon 3-7-45 expression. **A.** Replication profile of the left end of Chromosome III in cells harboring the GALp vector or *GALp-SLD3-SLD7-CDC45*. For details, see (Tanaka et al., 2011). **B.** Cells harboring *GALp-SLD3-SLD7-CDC45* were arrested in G1 and synchronously released into HU-containing media after induction of genes by addition of galactose (ON). Chromosomal DNA was isolated, separated using alkaline gels, and detected with *ARS305-* and *ARS302-320*-specific probes. **C.** *HMLα::URA3* and *HMLα::URA3* cells harboring *GALp* vector, *GALp-3-7-45, GALp-3-7, GALp-3-45, GALp-3-45^JET1^*, *GALp-7-45*, or *Δsir1* were serially diluted, spotted on the indicated plates, and grown at 30°C. **D.** *HMLα::URA3* harboring *GALp-3-7-45* (wt), its *Δswe1* and *Δmih1* derivatives, and *HMLα::URA3 Δsir1* cells were serially diluted, spotted on the indicated plates, and grown at 30°C. **E.** The cells used in D were grown in YPA-galactose, and their DNA contents were analyzed by flow cytometry.

**Supplementary Figure S2.**
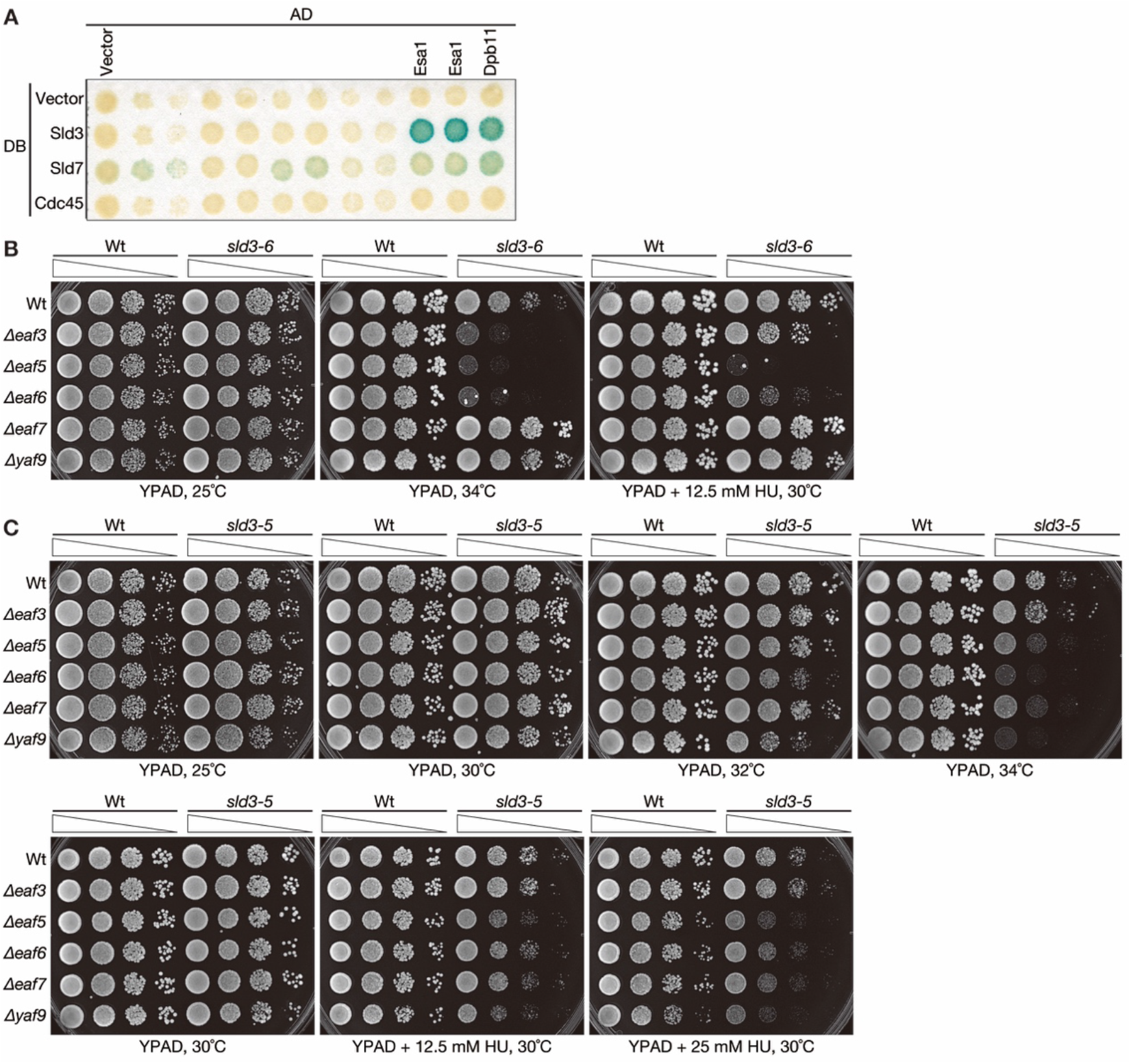
Sld3 interacts with Esa1. **A.** Original image of Fig. 2A. **B.** *Δeaf3, Δeaf5, Δeaf6, Δeaf7*, and *Δyaf9* cells in the wild-type (Wt) or *sld3-6* background were spotted and grown under the indicated conditions. **C.** *Δeaf3*, *Δeaf5*, *Δeaf6*, *Δeaf7*, and *Δyaf9* cells in the wild-type (Wt) or *sld3-5* background were spotted and grown under the indicated conditions.

**Supplementary Figure S3.**
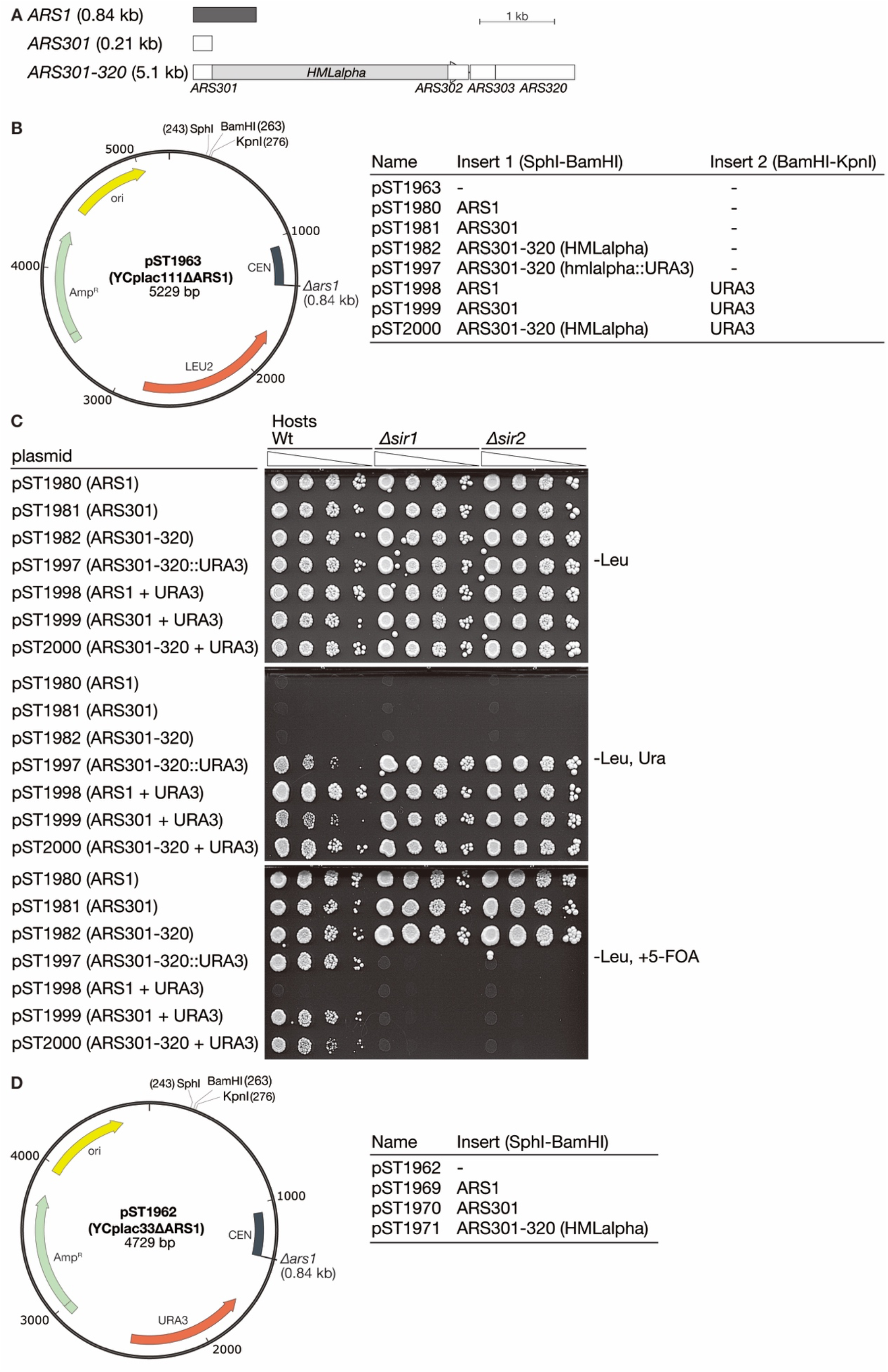
Structures of plasmids used in the plasmid loss assay. **A.** Schematics of the ARSs used. **B.** Schematics of the plasmid constructs used in C. pST1963 is a derivative of YCplac111 and lacks the *ARS1* sequence. *ARS1, HMLα ARSs* were inserted into the indicated site of pST1963 to yield pST1980-82. *URA3* was inserted into pST1980-82 to yield pST1997-2000. **C.** The indicated plasmids were introduced into the wild type (Wt), *Δsir1*, or *Δsir2*, and transformed cells were serially diluted, spotted, and grown on the indicated selective plates. **D.** Schematics of plasmids used in the plasmid loss assay in Figs. 3 and 4C.

**Supplementary Figure S4.**
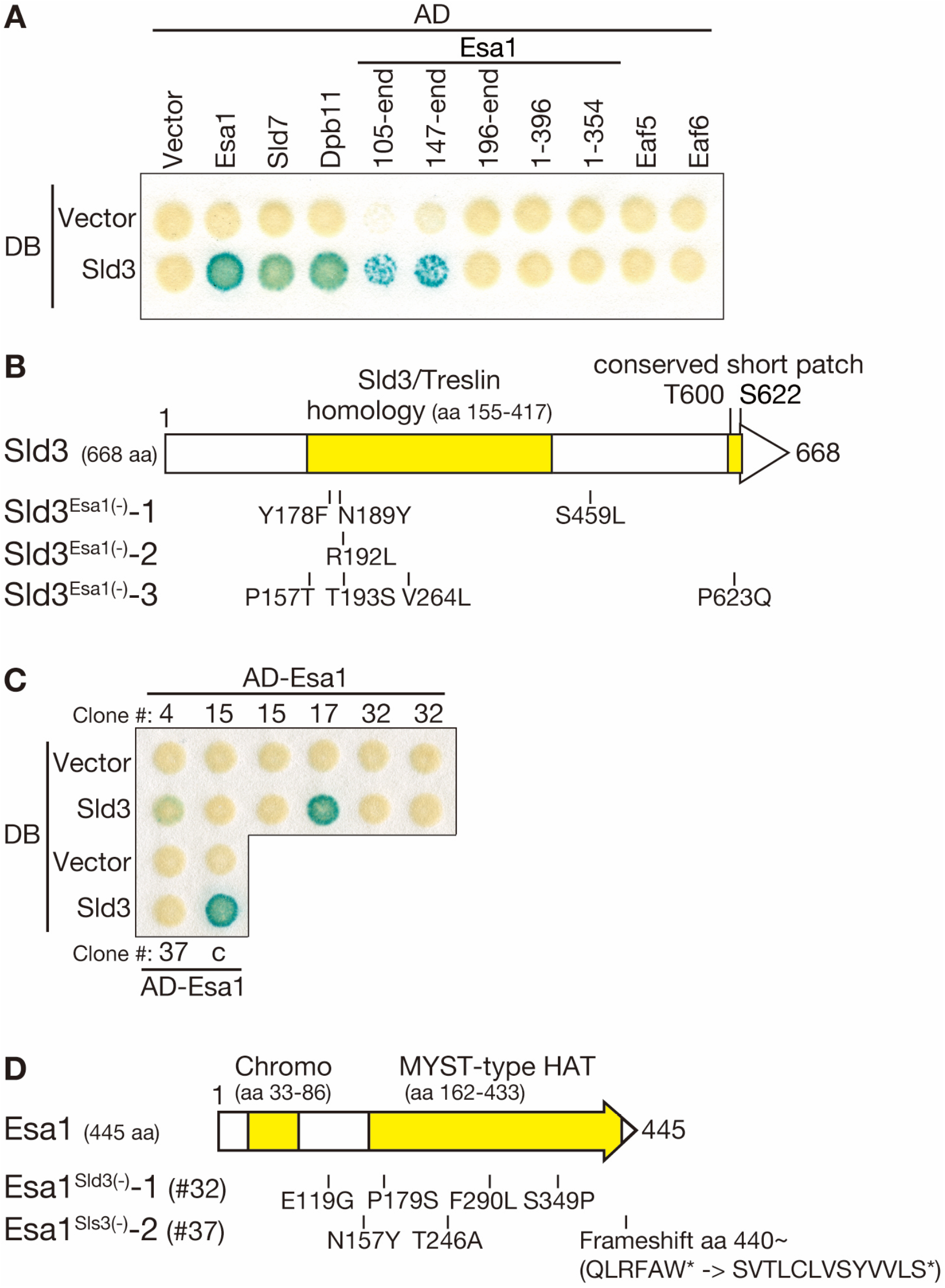
Isolation of mutants that diminish the Sld3–Esa1 interaction. **A.** Interaction between Sld3 and various Esa1 constructs, Eaf5, and Eaf6 were tested in the Y2H analysis. AD: activation domain. DB: DNA-binding domain. **B.** Schematics of Sld3^Esa1(-)^ mutant proteins. Numbers indicate the positions of amino acid residues. **C.** Isolation of Esa1^Sld3(-)^ mutants. Interactions between Esa1^Sld3(-)^ candidates and Sld3 were tested in the Y2H. Clone # indicates the initial number of candidates. #32 and #37 were named Esa1^Sld3(-)^-1 and −2, respectively. c: Positive control (Esa1) for the Y2H. Clone #15 lacked an interaction with Sld3 in this assay, as did #32 and #37. However, plasmid shuffling assay revealed that #15 could not support cell growth, whereas #32 and #37 could. Therefore, #15 was not retained as an Esa1^Sld3(-)^ mutant (see Materials and Methods). **D.** Schematics of Esa1^Sld3(-)^ mutant proteins. Numbers indicate the positions of amino acid residues.

**Supplementary Figure S5.**
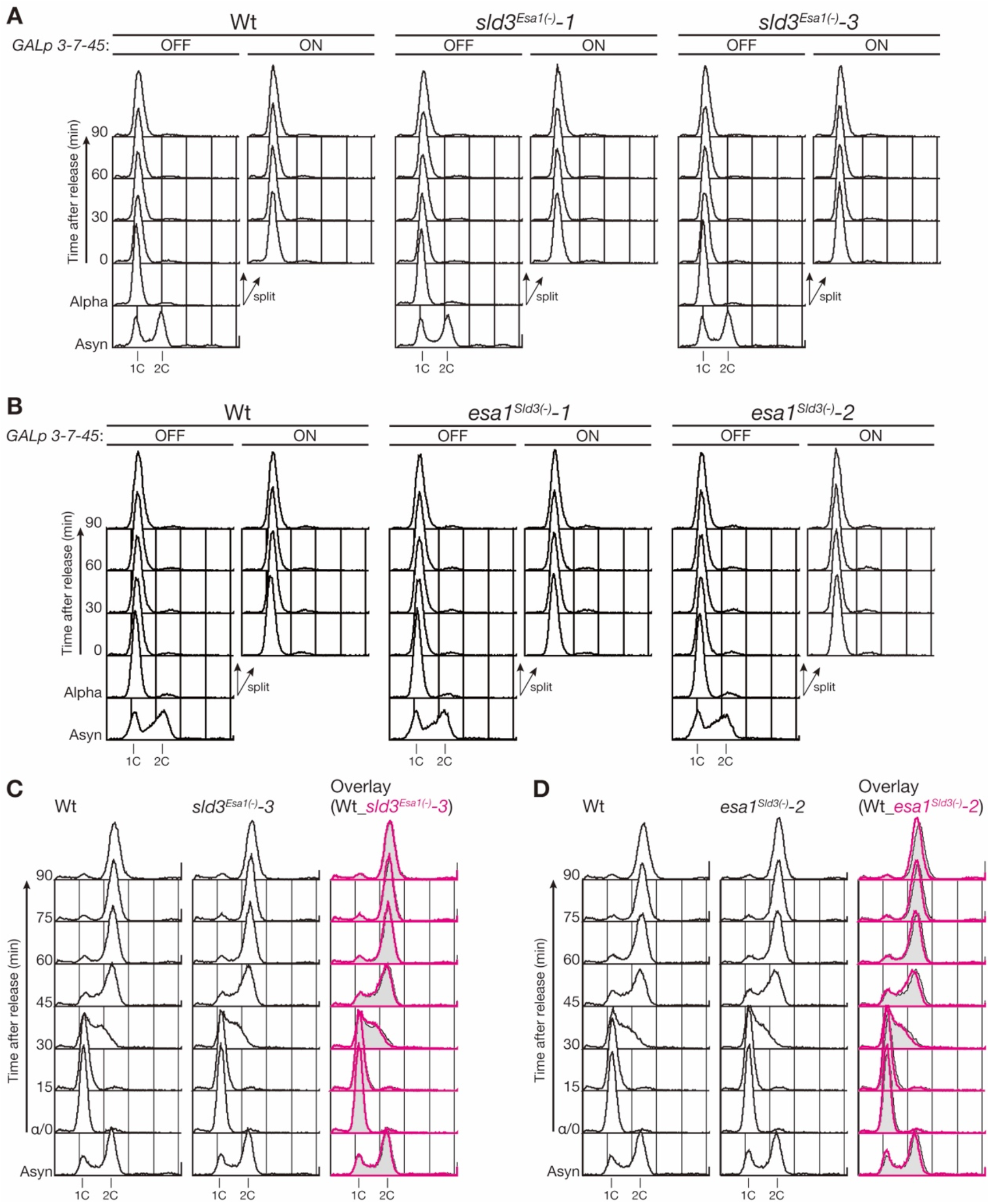
Analysis of the mutants that diminish the Sld3–Esa1 interaction. **A, B.** Aliquots of samples from Fig. 5A and 5B were analyzed by flow cytometry. **C, D.** Wild-type, *sld3^Esa1(-)^-3*, and *esa1^Sld3(-)^-2* cells were grown in YPAD at 25°C, arrested in G1 with α factor, and synchronously released. Samples were taken at the indicated times and analyzed by flow cytometry.

## Notes

### Competing Interest Statement

The authors have declared no competing interest.

